# Structural basis for allosteric regulation of human phosphofructokinase-1

**DOI:** 10.1101/2024.03.15.585110

**Authors:** Eric M. Lynch, Heather Hansen, Lauren Salay, Madison Cooper, Stepan Timr, Justin M. Kollman, Bradley A. Webb

## Abstract

Phosphofructokinase-1 (PFK1) catalyzes the rate-limiting step of glycolysis, committing glucose to conversion into cellular energy. PFK1 is highly regulated to respond to the changing energy needs of the cell. In bacteria, the structural basis of PFK1 regulation is a textbook example of allostery; molecular signals of low and high cellular energy promote transition between an active R-state and inactive T-state conformation, respectively. Little is known, however, about the structural basis for regulation of eukaryotic PFK1. Here, we determine structures of the human liver isoform of PFK1 (PFKL) in the R- and T-state by cryoEM, providing insight into eukaryotic PFK1 allosteric regulatory mechanisms. The T-state structure reveals conformational differences between the bacterial and eukaryotic enzyme, the mechanisms of allosteric inhibition by ATP binding at multiple sites, and an autoinhibitory role of the C-terminus in stabilizing the T-state. We also determine structures of PFKL filaments that define the mechanism of higher-order assembly and demonstrate that these structures are necessary for higher-order assembly of PFKL in cells.

## Introduction

Glycolysis is an ancient, highly-conserved metabolic pathway for the extraction of energy from sugars. During glycolysis, glucose is metabolized to produce energy in the form of ATP, the essential cofactor NADH, as well as other biosynthetic precursors to support cellular functions. The first committed step of glycolysis is catalyzed by phosphofructokinase-1 (PFK1), which converts fructose 6-phosphate (F6P) to fructose 1,6-bisphosphate (F1,6BP), consuming one molecule of ATP in the process. Given this central role as the “gatekeeper” of glycolysis, PFK1 is heavily regulated by the energy state of the cell; PFK1 is activated by signals of low cellular energy, such as AMP and ADP, and inhibited by signals of high cellular energy, such as ATP and citrate.

The structural basis for PFK1 regulation is best described for the bacterial enzyme^1–3^. Bacterial PFK1 is a D2-symmetric homotetramer with four active sites, each formed at an interface between two monomers. The enzyme transitions between an active R-state conformation, promoted by binding to F6P and allosteric activators, and an inactive T-state conformation, observed in the absence of F6P and upon binding to allosteric inhibitors. The R-state to T-state transition involves a rotation between dimers and rearrangement of active site residues, which together function to disrupt the F6P binding pocket^2^.

The PFK1 catalytic domain architecture is conserved in eukaryotes. However, eukaryotic PFK1 has an additional regulatory domain, which arose from gene duplication, tandem fusion, and evolution of the ancestral prokaryotic catalytic domain^4–6^. Generally, eukaryotic PFK1 tetramerizes via these regulatory domains, producing a quaternary structure distinct from that observed for the prokaryotic enzyme^7,8^. Oligomerization regulates PFKL activity; tetramers represent the active form of the enzyme, and allosteric activators promote tetramerization, while allosteric inhibitors promote tetramer disassembly^7,9–11^. Eukaryotic PFK1 is subject to additional layers of regulation, being regulated by over 20 allosteric ligands^12^, in addition to post-translational modification by phosphorylation, glycosylation, and acetylation^13–16^. Vertebrates possess three PFK1 isoforms: platelet (PFKP), muscle (PFKM), and liver (PFKL), each with particular catalytic and regulatory properties, as well as tissue-specific expression profiles^17^. PFKL forms filamentous polymers *in vitro* and micron-scale puncta in cells^18^. However, whether higher-order assembly of PFKL in cells reflects the filament formation observed with purified protein is unclear. Further, the functional role of these higher-order assemblies remains an open question.

Bacterial PFK1 provides a canonical example of the structural basis for allosteric regulation, though little is known about the structural basis for the regulation of the vertebrate enzyme. Existing crystal structures of vertebrate PFK1 in various ligand states all resemble the R-state, suggesting that crystal packing may preferentiallyselect for the R-state conformation^6–8^. Here, we determine cryoEM structures of PFKL in the R- and T-state conformations. The conformation of T-state PFKL differs from its bacterial counterpart and other vertebrate PFK1 structures. The T-state conformation of PFKL is stabilized by binding of the C-terminus across the catalytic and regulatory domains, and truncation of the C-terminus disrupts PFKL regulation. We further show that PFKL forms filaments in both the R- and T-state conformations, present cryoEM structures of filaments in both states, and demonstrate that micron-scale, punctate assemblies of PFKL observed in cells are composed of the same filament structures observed *in vitro*.

## Results

### Structure of human PFKL in the R-state

We determined structures of human PFKL in the presence of substrates ATP and F6P by cryoEM (Fig. 1). 2D classification produced an incompletely separated mixture of short filaments and free tetramers, which we therefore separated by 3D classification and refined independently (Fig. 1a,b, Fig. S1). Free tetramers and tetramers within filaments are in the same conformation (Cɑ RMSD 0.5 Å) (Fig. 1c), such that combining them in a masked, consensus refinement produced the highest resolution tetramer structure (Fig. 1d) (3.1 Å, 3.6 Å, and 2.9 Å resolution was achieved for the free tetramer, filament, and consensus tetramer structures, respectively; Fig. S1b). F6P and ADP are bound in the active sites (Fig. 1e), suggesting catalysis and product formation occurred during sample preparation. ADP is bound at two additional sites on each monomer: an activating allosteric site (site 1) present at the interface between the catalytic and regulatory domains–this site was previously shown to bind ADP, AMP, and a variety of small-molecule activators^6,19^ – as well as a second site (site 2) present in the catalytic domain, adjacent to loop 390-400 connecting to the regulatory domain (Fig. 1e). The allosteric sugar binding site is also occupied – we modeled this ligand as F1,6BP, likely produced by catalysis of F6P and ATP by PFKL (Fig. 1e). The F6P-bound PFKL structure presented here is in the active R-state conformation, as revealed by comparison to existing prokaryotic and eukaryotic R-state PFK structures (Fig. 1f)^3,6,8^.

**Fig. 1:**
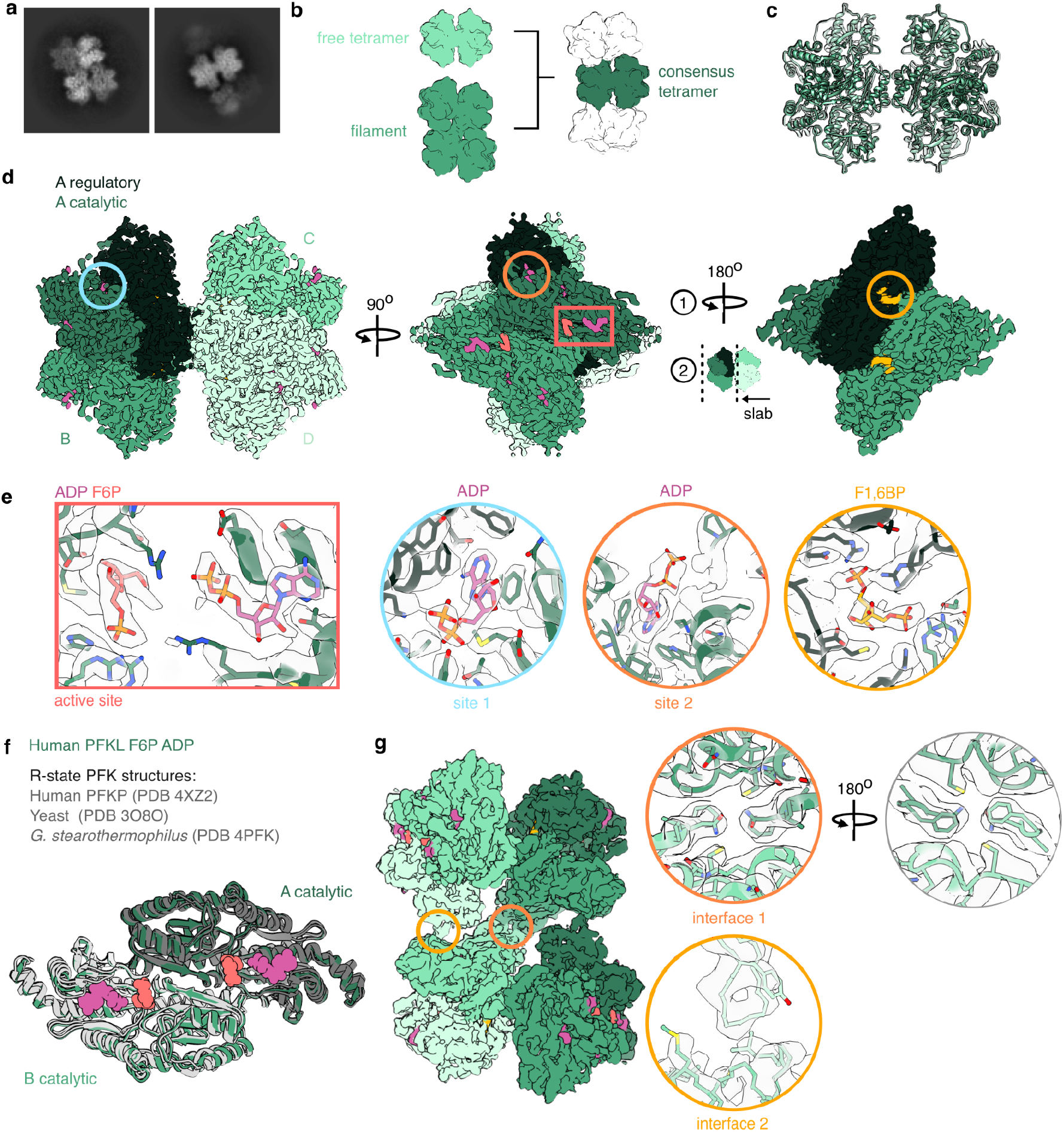
cryoEM structures of PFKL tetramers and filaments in the R-state. **a,** Representative 2D class averages of PFKL in the presence of F6P and ATP. **b,** Generalization of the cryoEM processing scheme. Three structures were determined: a free tetramer, a filament, and a consensus tetramer including both free and filament-associated tetramers. **c,** Overlay of the three structures outlined in **b**, which are in the same conformation. **d**, CryoEM structure of the consensus PFKL tetramer, colored by monomer. Monomer A is also colored by domain. **e,** Zoomed-in views of the regions indicated in **(d)** showing the various ligand-binding sites. **f,** F6P- and ADP-bound PFKL is in the R-state conformation, as revealed by comparison to existing R-state PFK structures. Catalytic domains are shown, with F6P and ADP colored as in panel (e). Structures are aligned on catalytic domain A. **g,** CryoEM structure of the R-state PFKL filament, colored by monomer. Circles show zoomed-in views of filament interfaces 1 and 2. Two views of interface 1 are shown, focused on N702 (left) and F700 (right) at the center of the interface.

PFKL filaments assemble as stacked tetramers via two separate interfaces (Fig. 1g), as previously described at lower resolution in a negative stain EM reconstruction^18^. Although PFKL filaments can adopt distinct straight or bent architectures^18^, adjoining tetramers are all related by 2-fold symmetry, and we therefore imposed C2 symmetry on tetramer pairs during 3D refinement (Fig. S1a). Interface 1, the largest interface, is on the symmetry axis and is formed by loops containing residues 511-520 and 693-705 of the regulatory domain. The core of the interface is formed by N702-N702 and F700-F700 interactions, with possible additional interactions between R695 and Y514. Interface 2 also involves residues on loop 693-705 of a different PFKL protomer, which insert into a groove on the surface of the catalytic domain of the adjoining tetramer. Masked local refinement of interface 1 improved the resolution from 3.6Å to 3.5Å (Fig. 1g, S1a) while two-fold symmetry expansion and local refinement of interface 2 failed to improve the EM map in this region. The precise interactions at interface 2 are therefore less well-resolved (Fig. 1g).

### Structure of human PFKL in the T-state

We next aimed to determine the structure of PFKL in the inactive T-state, and therefore solved cryoEM structures of PFKL in the presence of ATP but without substrate F6P (Fig. 2, Fig. S2). We also included F1,6BP in the sample preparation, as we suspected that occupying additional ligand sites might aid in stabilizing the protein for structure determination. We again observed a mixture of tetramers and short filaments by 2D classification, and therefore processed the data as described for the R-state structures; we again found that free tetramers and filament-associated tetramers were in the same conformation (Cɑ RMSD 0.39 Å), such that a consensus structure combining both achieved the highest resolution (2.9 Å, 3.1 Å, and 2.6 Å resolution was achieved for the free tetramer, filament, and consensus tetramer structures, respectively; Fig. S2b). On each monomer, ATP is bound in the active site, site 2, and a third site (site 3) between the catalytic and regulatory domains that does not contain a nucleotide in the R-state, while activating site 1 is not occupied (Fig. 2a). F1,6BP is bound in the allosteric sugar binding site (Fig. 2a).

**Fig. 2:**
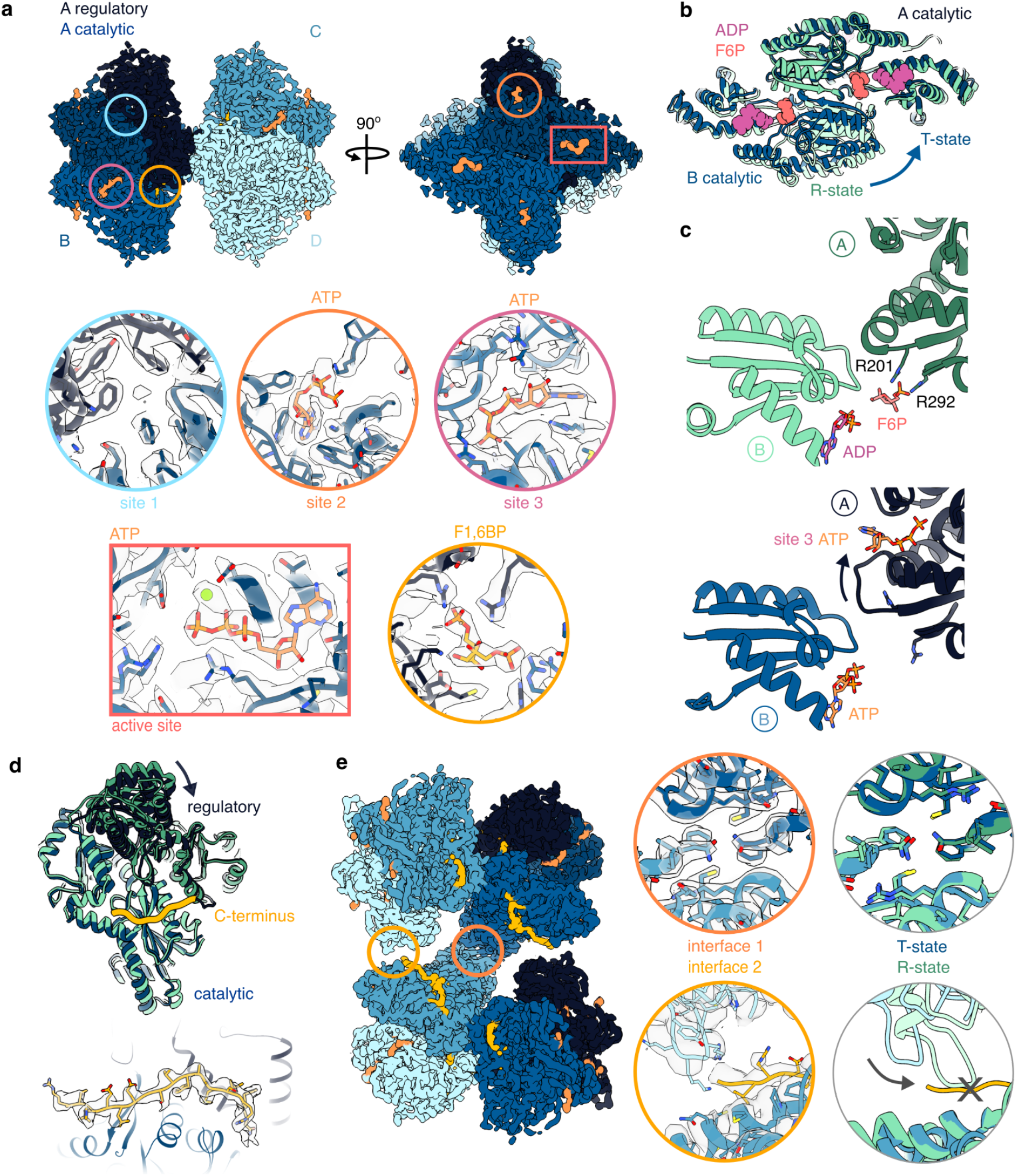
cryoEM structures of PFKL tetramers and filaments in the T-state. **a,** CryoEM structure of the consensus PFKL tetramer bound to ATP and F1,6BP, colored by monomer. Monomer A is also colored by domain. Insets show the various ligand binding sites. **b,** Comparison of the catalytic domain conformation of the ATP- and F1,6BP-bound structure (blue) with the R-state PFKL structure (green). Structures are aligned on the catalytic domain of subunit A. ADP and F6P from the R-state structure are shown in the active site. **c,** Comparison of the F6P binding site in the R-state (green) and T-state (blue) structures. The binding site involves residues from the catalytic domains of two monomers (A and B). **d,** Comparison of monomer conformations in the R-state (green) and T-state (blue) structures. Structures are aligned on their catalytic domains (lighter shades). The C-terminus (yellow) bridges across the regulatory and catalytic domains in the T-state conformation. Lower panel shows the isolated density of the C-terminus from the T-state cryoEM map. **e,** CryoEM structure of the T-state PFKL filament, colored by monomer. Circles show zoomed-in views of filament interfaces 1 and 2. Panels at right show comparisons to interfaces 1 and 2 in the R-state structure. Inward rotation of the catalytic domain at interface 2 in the R-state would produce a clash with the C-terminus at the position observed in the T-state structure.

In the absence of F6P, PFKL adopts the T-state conformation, characterized by a rotation between pairs of catalytic domains, which disrupts the F6P binding pockets without affecting ATP binding in the active sites (Fig. 2b). The PFKL T-state conformation differs from existing bacterial T-state PFK1 structures, exhibiting an additional rotation between adjoining catalytic domains, as well as structural rearrangements in the region around ATP sites 2 and 3 (Fig. S3a)^2,20^. In the PFKL structure, ATP binding at site 3 is linked to local unfolding of an adjacent ɑ-helix (Fig. 2c). This disrupts the positions of residues R201 and R292, which in the R-state bind to the phosphate of F6P in the active site of the adjoining catalytic domain (Fig. 2c). ATP binding in site 3 therefore appears to be inhibitory, contributing to disruption of the F6P binding pocket and stabilization of the T-state conformation. In existing R-state human PFKP structures, site 3 is occupied by a phosphate ion (Fig. S3b), and both phosphate and sulfate ions have been shown to increase PFK1 activity^21^. Taken together, this suggests that PFKL site 3 may function as both an allosteric activating and inhibitory site. Similarly, in bacterial PFK1 structures, site 3 is occupied by the allosteric inhibitor phosphoenolpyruvate in the T-state^20^ and the allosteric activator ADP in the R-state^3^, suggesting this site may also have a dual activating and inhibitory role in the bacterial enzyme (Fig. S3b).

Notably, rearrangement of active site residue R201 in PFKL differs from bacterial PFK1, where the corresponding residue R162 instead swaps positions with E161 upon F6P binding and transition to the R-state conformation (Fig. S3c). The T-state PFKL structure also differs from existing human PFKP crystal structures, which all exhibit overall catalytic domain conformations resembling the R-state, regardless of F6P occupancy^7,8^. Most of the PFKP structures also have an R-state arrangement of active site residues, with the exception of an ATP-bound structure (PDB 4XYJ) which exhibits the T-state orientation of R210 (R201 in PFKL) as well as local unfolding similar to that seen adjacent to ATP site 3 in T-state PFKL (Fig. S3d).

Transition from the R-state to T-state of PFKL also involves a rotation of the regulatory domain towards the catalytic domain within each monomer (Fig. 2d). The rotation compresses the activating site 1, preventing ADP/AMP binding at this site in the T-state conformation (Fig. 2a). Additionally, the C-terminal tail of PFKL (residues 754-771), which is disordered in the R-state structure, is bound in an extended conformation that bridges across the regulatory and catalytic domains in the T-state structure. The tail acts as a kind of strap linking the two domains, suggesting it may function to stabilize the inactive enzyme conformation (Fig. 2d).

T-state PFKL filaments have a similar overall architecture to R-state filaments, being composed of stacked tetramers (Fig. 2e). Interface 1 is the same in the T-state and R-state filament structures (Fig. 2e), while interdomain rotations in each protomer result in disruption of interface 2 interactions. In the T-state filament, loop 693-705 is rotated away from the binding groove on the adjoining tetramer, which is instead occupied by the C-terminus (Fig. 2d,e). Residues on loop 693-705 are in range to form contacts with the C-terminus in the T-state filament, though specific contacts were not well-resolved in this region. Inward rotation of loop 693-705 upon transition to the R-state conformation is sterically inhibited by a bound C-terminus in an adjoining T-state tetramer; in order for a tetramer within a filament to transition from the T-state to the R-state conformation, the C-termini of neighboring T-state tetramers must be displaced. Thus, competition at interface 2 between binding of a protomer’s own C-terminus and loop 693-705 on a neighboring tetramer may provide a mechanism to allosterically couple T- to R-state transitions between tetramers within filaments. For both the R- and T-state filaments, the two adjoining protomers at interface 2 exhibit the same overall conformation despite being in different packing environments, with one protomer donating and the other receiving loop 693-705 (Cɑ RMSD 0.5Å and 0.2Å for the R- and T-state structures, respectively) (Fig. S3e-g).

### Cellular assembly of PFKL requires filament formation

Towards understanding the function of PFKL filaments, we generated a non-polymerizing PFKL mutant. PFKP does not assemble filaments^18^,and several residues at PFKL interface 1 differ between the isoforms (Fig. 3a). N702 at the core of PFKL interface 1 is substituted for threonine in PFKP, and we therefore expressed and purified the substitution mutant PFKL-N702T (Fig. S4a). Negative stain EM confirmed that PFKL-N702T disrupted filament assembly in both R- and T-states, but did not disrupt tetramer formation (Fig. 3b). This result was confirmed by mass photometry: wild-type PFKL was predominantly filamentous, with 60% of protein in assemblies larger than tetramers, while PFKL-N702T was 90% tetrameric with no larger species observed (Fig. 3c). Both samples also exhibited small populations of monomers and dimers (Fig. 3c). Enzyme assays revealed that filament formation had only a modest effect on enzyme activity and allosteric regulation: Compared with wild-type PFKL, PFKL-N702T exhibited a slight increase in maximal velocity, F6P affinity and cooperativity, as well as a slight decrease in affinity for inhibitory ATP (Fig. S4b).

**Fig. 3:**
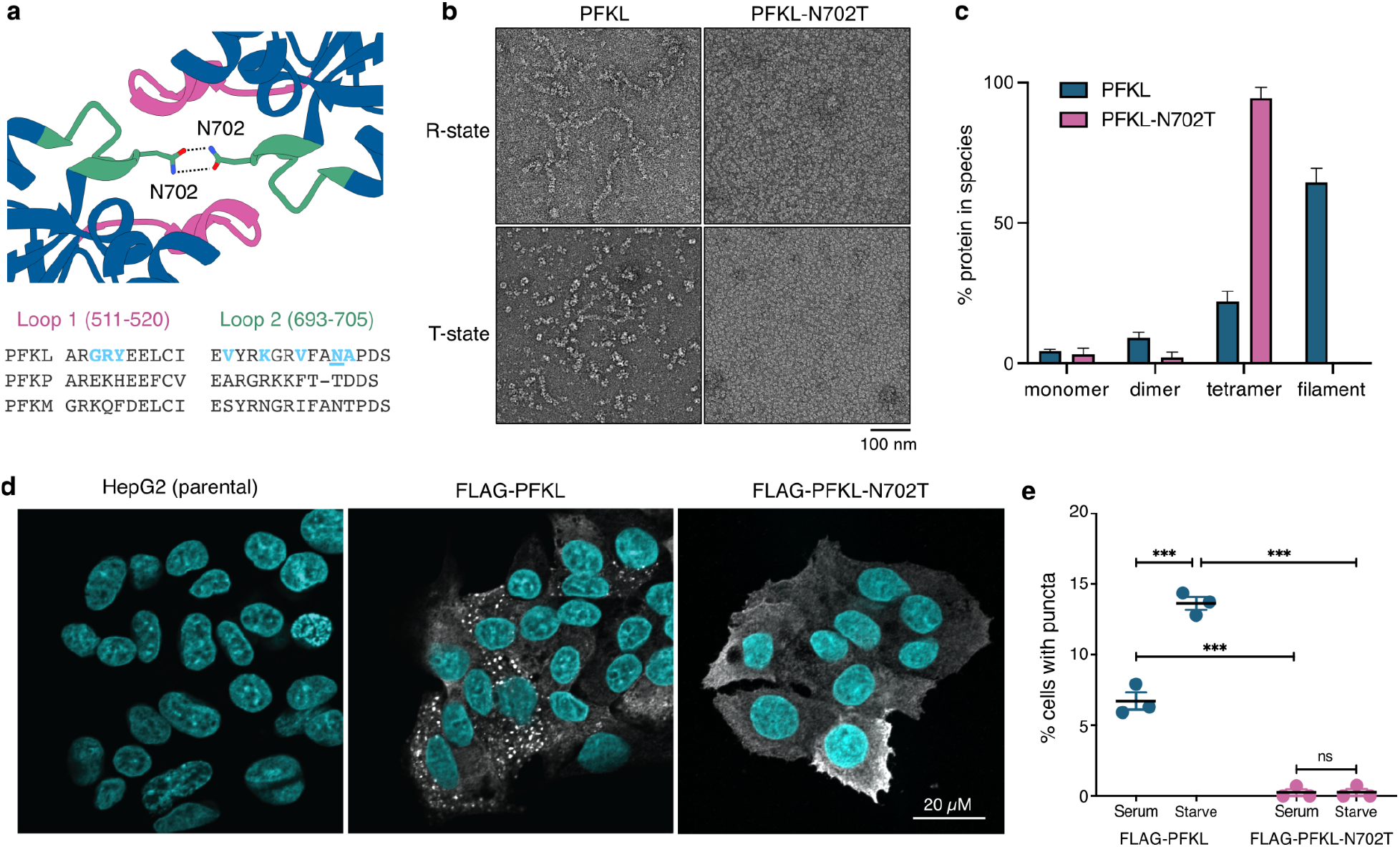
Filament interface 1 is essential for assembly of PFKL *in vitro* and in cells. **a,** Structure of PFKL filament interface 1 showing two loops forming the interface (top) and sequence alignment of interface 1 residues with other human isoforms (bottom). Poorly-conserved PFKL interface residues are highlighted in blue, with N702 underlined. **b,** Negative stain EM showing that PFKL-N702T does not form filaments in either R-state or T-state ligand conditions. Scale bar is 100 nm. **c,** Quantification of assembly state of PFKL and PFKL-N702T by mass photometry. **d,** Immunofluorescent images of parental HepG2 cells and cells expressing wild type FLAG-PFKL or FLAG-PFKL-N702T labeled with anti-FLAG (white) and Hoechst 33342 (blue). Puncta are only visible in wild type FLAG-PFKL cells. Scale bar is 20 µM. **e,** Quantification of the number of cells with PFKL puncta determined in continuous serum or in serum starved conditions. Data is representative of 3 independent passages of cells. Mean and SEM are shown. *** = P<0.001.

PFKL assembles into micrometer-sized structures in the human hepatocellular carcinoma cell line HepG2^22^. We aimed to determine whether these higher-order structures observed in cells correspond to the PFKL filaments observed with purified protein. We therefore expressed N-terminally-tagged wild-type FLAG-PFKL or non-polymerizing FLAG-PFKL-N702T in HepG2 cells and imaged both by immunofluorescence (Fig. 3d, Fig. S4c). In cells grown in complete media containing 10% FBS, FLAG-PFKL had a diffuse cytoplasmic localization in a majority of cells. However, in 7% of cells FLAG-PFKL assembled into multiple structures greater than 1 µm in diameter (Fig. 3d,e). We next asked if serum starvation altered the number of cells displaying puncta as withdrawal of insulin signaling decreases glucose uptake in HepG2 cells ^23^. In HepG2 cells that were incubated in media containing 0.1% FBS overnight, the percentage of cells containing FLAG-PFKL puncta doubled to 14% (Fig. 3d,e), suggesting that PFKL assembly is regulated by hormonal or growth factor signaling. In contrast, FLAG-PFKL-N702T was diffuse in both complete serum and serum starved conditions (Fig. 3d,e), indicating that polymerization of PFKL into filaments is required for assembly of the micron-scale PFKL structures observed in cells.

### The T-state of PFKL is stabilized by the C-terminal tail

The C-terminal tail is required for proper regulation of PFK1 in *Dictyostelium discoideum* and human PFKP^8,24^, though the mechanism by which it imparts allosteric control of PFK1 has not been described. Notably, the C-terminal region of PFK1 shows substantial sequence variation across species and also between the PFKL, PFKP, and PFKM isoforms (Fig. S4d). To determine the role of the C-terminal tail in PFKL allosteric regulation, we purified a construct in which the C-terminal 28 amino acids were removed (PFKL-ΔC) and tested its activity and regulation. As shown for other human PFK1 isoforms^8^, removal of the C-terminal tail increased affinity for the sugar substrate F6P and relieved allosteric inhibition by ATP (Fig. 4a, Table 1). Negative stain EM and mass photometry indicated that truncation of the C-terminus had no major effect on tetramer or filament formation compared with the wild type enzyme (Fig. S4e,f).

**Fig 4:**
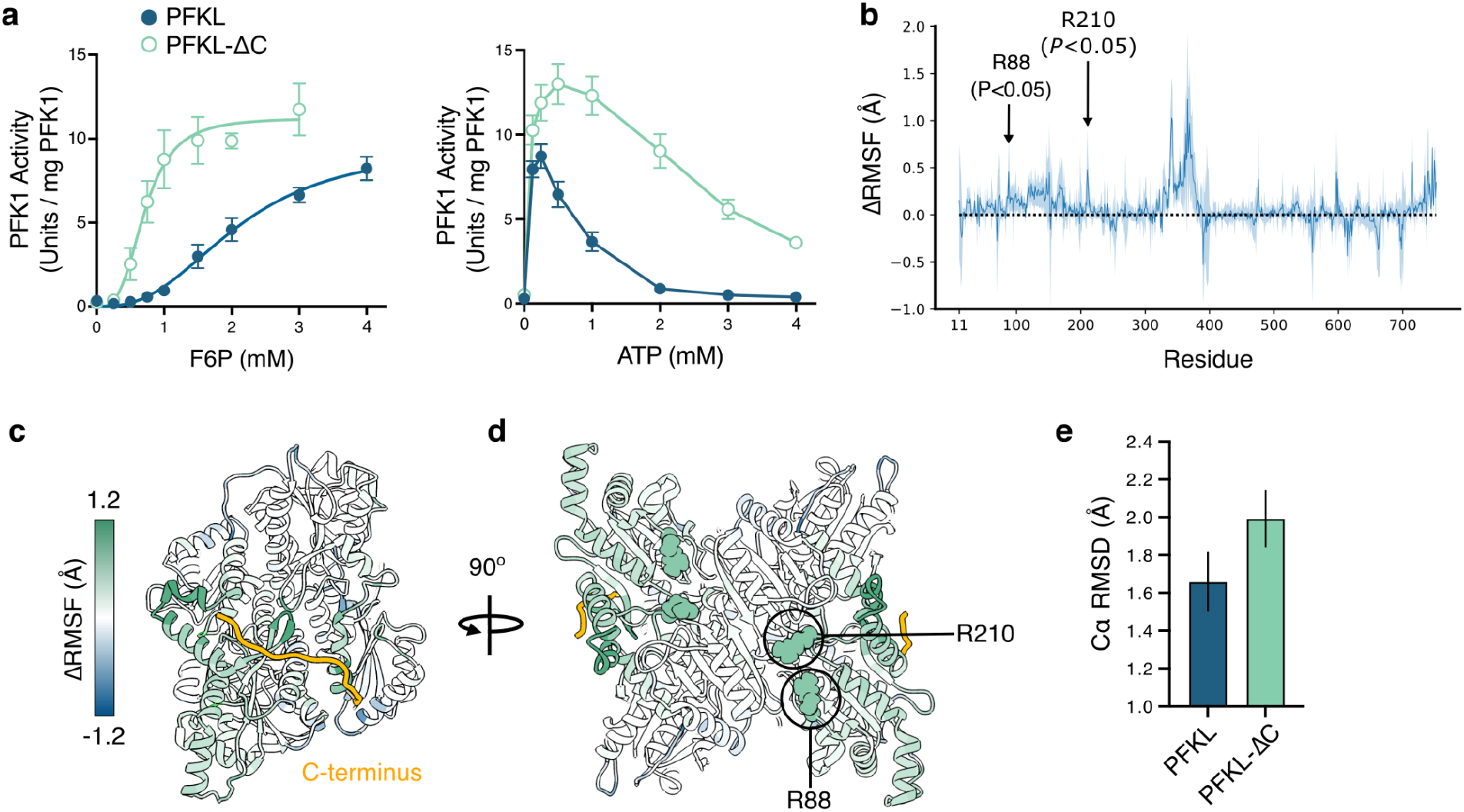
The C-terminal tail of PFKL stabilizes the T-state conformation. **a,** Enzyme activity assays to determine the affinity for F6P (left) and ATP (right) for wild type PFKL (blue) and PFKL-ΔC (green). Assay conditions are described in Table 1. Data are mean and SEM. **b,** Difference in per-residue root-mean-square fluctuations (ΔRMSF) after C-terminal tail removal as observed in MD simulations. The light blue regions correspond to the 95% confidence intervals. Arginine residues 88 and 210, involved in binding F6P in the R-state, are highlighted by arrows. **c,d,** PFKL monomer with residues colored according to ΔRMSF values in **(b)**. Removal of the C-terminus produces elevated ΔRMSF values at the C-terminus binding site **(c)** and in the active site **(d)**. **e,** Average Cα RMSD from the cryoEM structure of a T-state subunit during the last 200 ns of the MD simulations for PFKL and PFKL-ΔC. Error bars represent the 95% confidence intervals.

**Table 1:** Kinetic parameters for wild type and mutant PFKL.

| | $K_{m,F6P} \pm \text{SEM}$<br>(mM) | Hill coefficient $\pm \text{SEM}$ | $V_{\text{max}} \pm \text{SEM}$<br>(Units per mg PFK1) |
| --- | --- | --- | --- |
| PFKL (n=7) | $2.1 \pm 0.3$ | $2.6 \pm 0.6$ | $9.6 \pm 1.4$ |
| PFKL-N702T (n=6) | $1.5 \pm 0.1$ | $3.9 \pm 0.7$ | $13.8 \pm 0.8$ |
| PFKL-$\Delta C$ (n=5) | $0.8 \pm 0.1$ | $2.6 \pm 0.5$ | $10.6 \pm 0.7$ |

The kinetic parameters for PFKL were determined at pH 7.5 with 125 $\mu\text{M}$ ATP. The parameters were generated by fitting data to the allosteric sigmoidal model. The number of experimental replicates is noted in brackets.

Given that the PFKL C-terminal tail was only resolved in the T-state cryoEM structure, we hypothesize that the C-terminal tail stabilizes PFKL in the less active T-state conformation and removal of these residues would promote transition to the R-state. To test this hypothesis, we performed molecular dynamics (MD) simulations and asked if removal of the tail altered the conformation of the enzyme. We ran three independent 400 ns MD simulations with the C-terminal tail included (residues 753-780) using the CHARMM36m protein force field^25^, and we repeated the same computational protocol for the structure lacking the C-terminal tail. The simulations showed that removal of the C-terminus destabilizes the T-state structure, as quantified by the difference in per-residue root-mean-square fluctuations (ΔRMSF) (Fig. 4b-d). The structural changes mainly concerned the region of residues ∼330-380 and notably residue 715 on the solvent-exposed side of the subunit. These residues correspond to the region of PFKL where the C-terminal tail binds. Additionally, some of the residues surrounding the catalytic site show a change in structure, specifically residues involved in binding F6P (R88 and R210), suggesting that conformational changes caused by removal of the C-terminal tail propagate to the active site. The overall conformation of PFKL subunits transitions away from the T-state upon the C-terminal tail removal, as measured by the average Cα RMSD from the cryoEM structure of the T-state (Fig. 4e).

## Discussion

The regulation of PFK1 is a textbook example of allostery, with substrate, activators, and inhibitors competing to shift the enzyme between the active R-state and inactive T-state conformations. However, much of the structural insight into PFK1 regulation has been limited to the bacterial enzyme, with all existing structures of vertebrate PFK1 exhibiting the R-state conformation^7,8^. The R-state PFKL structure presented here resembles existing R-state PFK1 structures indicating–perhaps unsurprisingly–that a particular, conserved conformation is necessary for catalytic activity. By contrast, T-state PFKL differs from existing T-state structures, exhibiting a different overall conformation, an additional allosteric ATP-binding site, as well as additional regulation through binding of the C-terminal tail. Previous studies have shown that insulin induces phosphorylation of the PFKL C-terminus, increasing PFKL activity and flow through glycolysis^26^. Based on the structures presented here, it seems plausible that phosphorylation could displace the PFKL C-terminus, thus favoring the R-state conformation and increasing enzymatic activity. There is substantial sequence variation in the C-terminal tails between human PFKL, PFKP, and PFKM, perhaps providing an additional mechanism for differential regulation of the isoforms. Given the large number of ligands known to regulate PFK1 activity, additional structures in different ligand states are certainly warranted, and may lead to identification of additional PFK1 conformations.

PFKL filaments are somewhat unique in comparison to filaments of other metabolic enzymes. As with many other enzyme filaments, PFKL filaments are polymers of a subunit with dihedral symmetry^27–29^ – filaments of this sort may be the simplest to evolve, with a single, symmetric interface allowing for propagation of an unbounded polymer ^30,31^. For most enzyme filaments, the helical axis is coincident with a symmetry axis within the core, dihedral protomer, producing filaments with a linear architecture. With PFKL filaments, however, incoming tetramers can associate at one of two different positions on a filament end, resulting in both straight and bent architectures^18^. Notably, these variable architectures emerge despite the fact that each individual filament interface is the same.

Enzyme filaments variously function to increase or decrease enzyme activity^32–35^, increase cooperativity in regulation^36^, facilitate substrate channeling^37,38^, and tune affinity for regulatory ligands^39–42^. PFKL filaments differ in having no major direct effect on enzyme activity or regulation, and their function may therefore only be apparent in the context of the cell. Indeed, under conditions of energy stress, PFK1 assembles with other glycolytic enzymes into cellular foci which appear to both increase the rate of glycolysis and localize energy production to particular subcellular sites^22,43,44^. In neurons, for example, PFK1 assemblies appear at synapses, where there is an increased demand for ATP production^43^. PFKL assemblies have also been observed near the plasma membrane, where they could function to increase local ATP concentrations to support endocytosis, protrusion, and active transport^18^. These assemblies may increase the rate of glycolysis by enabling more efficient transfer of substrates between members of the glycolytic pathway. The fact that PFKL can form filaments with a range of bent and straight architectures suggests that long-range order of PFKL oligomers may be less important than aggregation of many copies of the enzyme in one location in the cell. This would be consistent with a scaffolding role of PFKL in assembly of condensates containing multiple glycolytic enzymes. The PFKL-N702 non-polymerizing mutant provides a tool to address the functional role of PFKL assembly in cells, and to determine if the higher-order assembly of other glycolytic enzymes depends on PFKL polymerization.

In summary, the present study provides a detailed structural understanding of the allosteric regulation of eukaryotic PFK1. These structural insights will aid in the development of small molecule compounds specifically designed to target PFK1. Pharmacologically targeting glycolytic enzymes is a proposed therapeutic approach for the treatment of diseases in which glucose metabolism is dysregulated. For example, proliferative cancer cells have an increased reliance on glycolysis to generate energy and biosynthetic precursors to support their rapid proliferation, which may make them susceptible to inhibition of glycolytic flux. However, there are currently no specific, high-affinity PFK1 inhibitors available; generating small molecules that can stabilize the T-state, possibly by stabilizing the C-terminal tail-bound conformation, is an area of future research.

## Materials and Methods

### Expression, purification, and activity of recombinant PFKL

Recombinant human PFKL (NP_002617) with an amino terminal His-tag was generated as described^18^. Site-directed mutagenesis of PFKL was achieved through Gibson Assembly. The presence of the mutations and the integrity of the constructs were verified by Sanger sequencing. PFK1 activity was determined using an auxiliary enzyme assay to monitor the production of fructose 6-phosphate adapted for use in a 96 well format, as previously described ^7,18,45^. The absorbance at 340 nm was measured using a Varioskan LUX Multimode microplate reader (ThermoFisher). Kinetic parameters were generated by nonlinear regression analysis using Prism (GraphPad Software) and are the mean of a minimum of three measurements from two independent preparations of protein. One unit of activity was defined as the amount of enzyme that catalyzed the formation of 1μmol of fructose 1,6-bisphosphate per minute at 25°C. ATP inhibition was calculated at pH 7.5 with 4 mM F6P.

### Negative stain electron microscopy

Negative stain samples were prepared by diluting freshly thawed protein to 1 μM in 50 mM HEPES pH 7.5, 20 mM (NH_4_)_2_SO_4_, 10 mM MgCl_2_, 2 mM ADP, 2 mM F6P to promote the R-state conformation or in 50 mM HEPES pH 7.5, 20 mM (NH_4_)_2_SO_4_, 10 mM MgCl_2_, 4 mM ATP, 0.5 mM F1,6BP to promote the T-state conformation. The protein was applied to glow-discharged carbon-coated grids, washed three times with water, and stained with 2% (w/v) uranyl formate. Grids were imaged using a 120 kV FEI Spirit microscope with a 4k x 4k Gatan Ultrascan CCD camera.

### CryoEM sample preparation, data collection, and data processing

CryoEM samples were prepared by applying protein to CFLAT 2/2 holey carbon grids (Protochips) and blotting away liquid four times successively before plunging into liquid ethane using a Vitrobot (ThermoFisher). Samples used to determine R-state structures contained 9 µM PFKL, 2 mM F6P and 1 mM ATP in storage buffer (20 mM HEPES pH 7.5, 1 mM DTT, 500 µM ammonium sulfate, 5% glycerol, 100 µM EDTA). Samples used to determine T-state structures contained 2 mM ATP, 10 mM MgCl_2_, and 500 µM F1,6BP in storage buffer. For R-state PFKL structures, movies were collected on a Titan Krios (ThermoFisher) equipped with a K3 Direct Detect Camera (Gatan Inc.) operating in superresolution mode with a pixel size of 0.4215 Å/pixel, 75 frames, and a total dose of 63 electrons/Å^2^. For T-state PFKL structures, movies were collected on a Titan Krios equipped with a K2 Direct Detect Camera (Gatan Inc.) operating in superresolution mode with a pixel size of 0.525 Å/pixel, 50 frames, and a total dose of 90 electrons/Å^2^. A subset of the data for the T-state sample was collected at 30° tilt. Data collection was automated with Leginon software^46^. CryoEM data was processed using cryoSPARC^47^, and processing workflows are summarized in figures S1 and S2. Superresolution movies were motion corrected, dose-weighted, and binned 2X using patch motion correction. CTF parameters were estimated using patch CTF. Particles were picked using the blob picker and subjected to multiple rounds of 2D classification. Initial 3D refinements were performed with combined particles representing both filaments and free tetramers, using a tetramer starting reference. Tetramers and filaments were then separated by 3D classification (without alignment) and refined independently with D2 and C2 symmetry, respectively. Additional rounds of 3D classification were performed to remove poor quality and damaged particles (Fig. S1 and S2). Masked, local refinement of filament interface 1 was performed using a mass encompassing the two dimers associated across interface 1. Per-particle defocus, beamtilt, and spherical aberration were also refined for all structures. Density modification was performed using ResolveCryoem in Phenix^48^. Atomic models were built using ISOLDE^49^ and real-space refinement in Phenix. Structures were visualized with UCSF Chimera^50^. Cryo-EM data collection, refinement and validation statistics are summarized in table S1.

### Mass photometry

Mass photometry experiments were performed using the TwoMP system by Refeyn^51^. PFKL filaments were assembled by diluting freshly thawed PFKL to 2.5 μM in cold dilution buffer (50 mM HEPES pH 7.5, 100 mM KCl). The 2.5 μM PFKL sample was diluted to 250 nM in cold polymerization buffer (50 mM HEPES pH 7.5, 100 mM KCl, 20 mM (NH_4_)_2_SO_4_, 10 mM MgCl_2_, 2 mM ADP, 2 mM F6P). The samples were incubated on ice for 2 min before they were diluted 10-fold into a droplet of room temperature MP buffer (50 mM HEPES, pH 7.5, 50 mM KCl). As PFKL-N702T was predominantly tetrameric with no protein sequestered into larger order oligomers, it was diluted 20-fold for a final concentration of 12.5 nM to obtain a measurable number of binding events. Movies were recorded immediately following dilution. Binding events were measured at a rate of 600 frames/min for one minute. Mass calibrations were performed with 5-10 nM each of beta-amylase, apoferritin, and glutamate dehydrogenase. Movies were collected with AcquireMP v2.3 and data was analyzed using DiscoverMP v2.3.To compare the relative populations of PFKL species in each movie, the counts per species was determined by the area under the curve of a gaussian fit to the histogram of mass species using the DiscoverMP software. This number was normalized to account for the number of PFKL monomers per species and converted to a percent of the total counts per species per movie. These percentages were averaged across replicates and compared among PFKL wild-type and mutants. At least three replicates were performed for each sample.

### Generation and imaging of HepG2 cells

HepG2 cells were purchased from ATCC and were verified to be mycoplasma-free. Cells were grown in high glucose DMEM (ThermoFisher Scientific) supplemented with 10% Heat Inactivated Fetal Bovine Serum (FBS; ThermoFisher Scientific) and 1% penicillin/streptomycin (ThermoFisher. Scientific) in a 37°C tissue culture incubator with 5% CO_2_. FLAG-PFKL, and FLAG-PFKL-N702T were cloned into a pLenti6.3 vector (ThermoFisher Scientific). The presence of the mutations and the integrity of the constructs were verified by Sanger sequencing. HepG2 cells expressing FLAG-PFKL were generated as previously described^52^. Lentivirus was generated using HEK293^FT^ cells (ATCC) and ViralPower lentiviral packaging mix (ThermoFisher Scientific), according to the manufacturer instructions. Blasticidin-resistant cells were pooled and used in experiments. The expression of exogenous FLAG-PFKL was verified by western blot analysis using primary antibodies generated against FLAG (Millipore Sigma) while GAPDH antibodies (Cell Signaling Technology) were used as a loading control.

For immunofluorescent analysis, parental HepG2 cells or cells expressing FLAG-PFKL or FLAG-PFKL-N702T were plated on 1.5mm glass coverslips and incubated overnight at 37°C. Cells were washed with PBS and the media was replaced with media containing either 0.1% or 10% FBS. The next day, cells were rinsed in PBS, fixed in 4% paraformaldehyde, and permeabilized in 0.1% Triton X-100. Coverslips were blocked in blocking buffer (3% BSA, 1% non-immune goat serum, 1% cold water fish gelatin) at room temperature for 30 min. Primary antibodies to FLAG were used at a 1:1000 in blocking buffer and incubated at 37°C for 2 hours. Coverslips were incubated with secondary goat anti-mouse antibodies labeled with Alexa Fluor-488 and Hoechst-33342 for 30 minutes at 37°C. Coverslips were mounted on glass slides with Prolong™ Gold antifade reagent (Thermofisher Scientific), allowed to cure overnight, and sealed with nail polish. Coverslips were imaged using a Nikon TI2-E Inverted Microscope with a CSU-W1 Confocal Scanner spinning disk with a 60x oil immersion objective. Image processing was performed in NIS Elements and ImageJ. Approximately 100-150 cells were counted in each experimental group for three independent passages of cells. Cells were considered to contain PFKL assemblies if they contained more than three FLAG-PFKL punctae at least 1µm in diameter. Two-way ANOVA was performed to determine significance using GraphPad Prism.

### MD simulations

The cryoEM structure of the PFKL tetramer in the T-state including the ATP and F1,6BP ligands was used as the starting point for all-atom MD simulations. For simulations involving the full-length C-terminal tail, the initial positions of the nine C-terminal residues, which were not resolved in the cryo-EM structure, were modeled using PyMOL software version 2.5.4^53^. For simulations of the PFKL-ΔC construct, the structure of each subunit was truncated after residue 752. The first 10 N-terminal residues, which were missing from the experimental structure, were not included in the simulations. The protonation states of histidine residues were adopted from the experimental structure.

The GROMACS pdb2gmx tool^54^ (version 2022.5) was used for protein topology preparation, with the CHARMM36m^25^ force field being employed for the protein. Existing CHARMM parameters for ATP were obtained from the CHARMM-GUI webserver^55,56^, and the same web server was employed to parameterize the FBP molecule using the CHARMM General Force Field (CGenFF)^57^. Water molecules were described with the TIP3P model^58^, and K^+^ and Cl^-^ ions were modeled using the default ion parameters associated with the CHARMM36m force field. To allow for a longer time step in production simulations, the hydrogen mass repartitioning scheme^59^, transferring the mass of 3 Da to a hydrogen atom from the nearest heavy atom, was applied to protein and ligand hydrogen atoms.

The PFKL tetramer was placed in a rhombic dodecahedral simulation box with the nearest-image distance of 20.1 nm and 18.7 nm for the system with- and without the C-terminal tail, respectively. The protein was solvated in a 150 mM KCl aqueous solution, including additional K^+^ ions to neutralize the net negative charge of the ligand-bound protein. In total, the system contained ∼569,000 and ∼451,000 atoms, respectively.

All-atom MD simulations were performed in the GROMACS 2022.5 software^54^. Short-range non-bonded interactions were treated with a 1.2 nm cutoff, and van der Waals forces were smoothly switched to zero between 1.0 and 1.2 nm. Long-range electrostatic interactions were computed using the particle mesh Ewald method^60^. The leap-frog algorithm^61^ was used to numerically integrate Newton’s equations of motion. The LINCS algorithm^62^ was employed to constrain the lengths of all hydrogen-involving protein bonds, with the highest order in the expansion of the constraint coupling matrix being increased from 4 to 6 in production simulations to achieve better accuracy with the longer time step used. The SETTLE algorithm^63^ was utilized to keep water molecules rigid.

To gently relax the protein structure in the simulation box, the system was subjected to an energy-minimization and equilibration protocol. First, an energy minimization was performed to decrease the maximum force below 1000 kJ mol^−1^ nm^−1^, while harmonic restraints were applied to all heavy atoms of the protein and those of the ligands (with a force constant k_BB_ = 1000 kJ mol^−1^ nm^−2^ assigned to the protein backbone atoms as well as the ligand atoms and a force constant k_SC_ = 500 kJ mol^−1^ nm^−2^ applied to the side-chain atoms). This energy minimization was followed by a six-step relaxation protocol during which the harmonic position restraints were gradually decreased. The first two steps of the equilibration protocol were performed in the NVT ensemble and were followed by four NPT simulations (see Table S2 for more details).

The production simulations were performed with an extended time step of 4 fs, enabled by the use of the hydrogen mass repartitioning approach^59^. The length of each production trajectory was 400 ns. For each system, a set of three independent production trajectories was obtained, each started after independent minimization and equilibration of the initial structure. The trajectory frames were saved every 200 ps.

In equilibration simulations, the temperature of the system was maintained at 300 K by the Berendsen thermostat^64^ with a time constant of 1 ps, and where applicable, the pressure was kept constant at 1.01 bar by the Berendsen barostat with the same time parameter. In production simulations, performed at the same temperature and pressure as the equilibration, the velocity rescaling thermostat with a stochastic term^65^ was used with a time constant of 0.1 ps, and the Parrinello-Rahman barostat^66^ was employed with a time constant of 1 ps. In all cases, the thermostats were coupled separately to the protein and to the rest of the system.

The per-residue root-mean-square fluctuations (RMSF) were calculated separately for each subunit and over each production trajectory using the GROMACS RMSF tool^54^ by considering the positions of all the atoms forming each residue. To describe the effect of the C-terminal tail removal, the differences in the per-residue means of these values were calculated between the PFKL and PFKL-ΔC systems. The *P*-values as well as the 95% confidence intervals of these differences were determined by performing a two-sided unpaired *t*-test using the Pingouin package^67^.

The root-mean-square deviation of Cα atoms (Cα RMSD) relative to the cryoEM structure of the T-state was computed for residues 20–752 in each subunit after aligning the subunit with a cryoEM subunit. The Cα atoms of residues 20–752 were used for the alignment. For each subunit, the average Cα RMSD over the second half of the production trajectory was calculated, and the mean of these values obtained for all the subunits and independent production trajectories was reported together with 95% confidence intervals estimated using Student’s *t*-distribution.

## Data Availability

Cryo-EM structures and atomic models have been deposited in the Electron Microscopy Data Bank (EMDB) and Protein Data Bank (PDB), respectively, with the following accession codes: EMD-43747, PDB: 8W2G (R-state PFKL tetramer); EMD-43749, PDB: 8W2I (R-state PFKL filament); EMD-43748, PDB: 8W2H (T-state PFKL tetramer); EMD-43750, PDB: 8W2J (T-state PFKL filament).

## Acknowledgements

We are grateful to the Arnold and Mabel Beckman Cryo-EM Center at the University of Washington for the use of electron microscopes. This work was supported by the National Institutes of Health (1R35GM149542 and S10OD023476 to J.M.K.). S.T. was supported by the Czech Science Foundation (grant no. 23-06437S). B.A.W was supported by West Virginia University Start-up funding and Visual Sciences CoBRE project leader funding (P20GM144230).

---

**Fig. S1:**
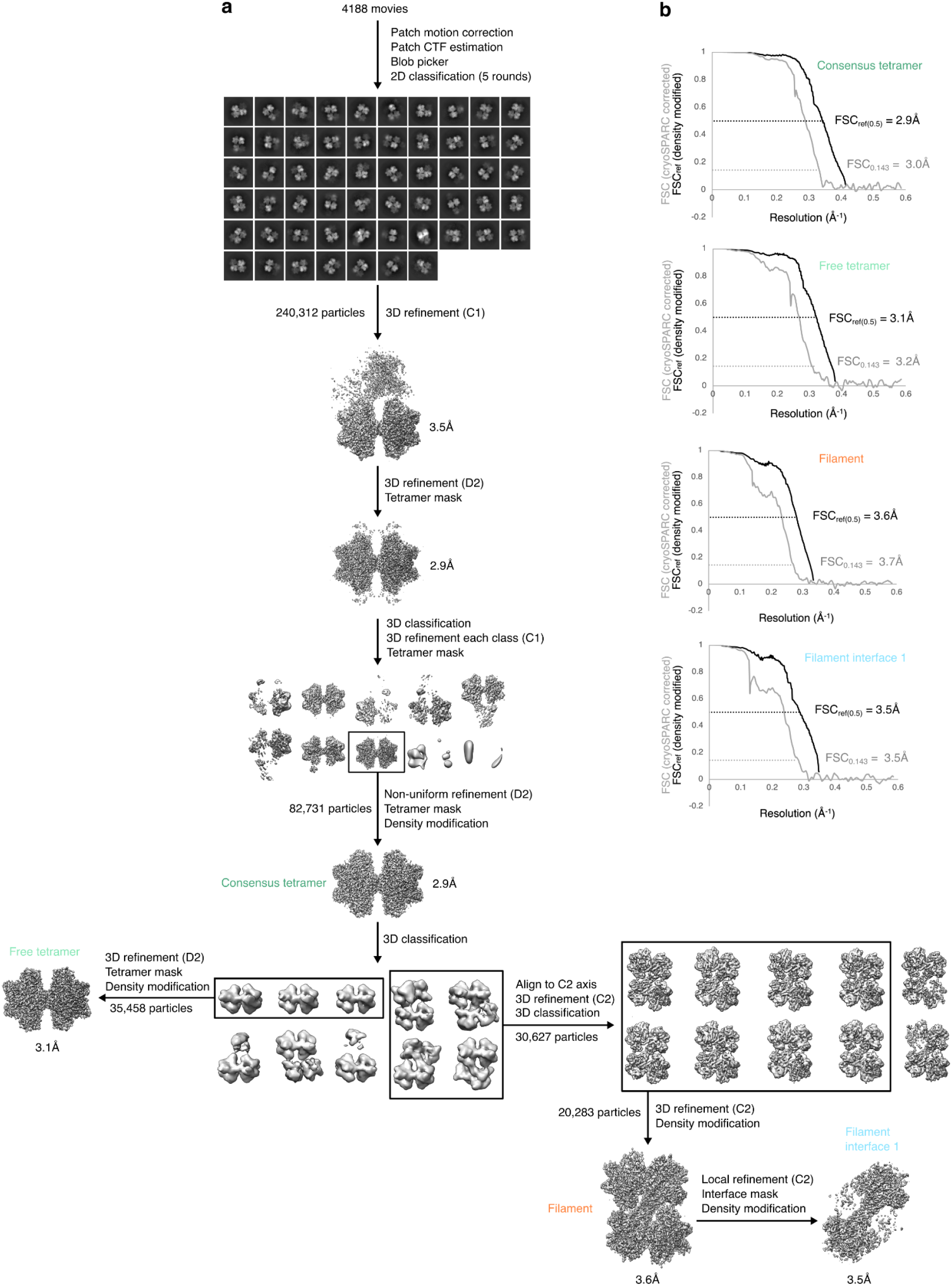
cryoEM data processing for R-state PFKL. **a,** Flowchart of cryoEM data processing. **b,** Noise-substituted corrected FSC curves (grey lines) and FSCref curves after density modification (black lines) and corresponding resolution estimates for the R-state consensus tetramer, free tetramer, filament, and filament interface 1 structures.

**Fig. S2:**
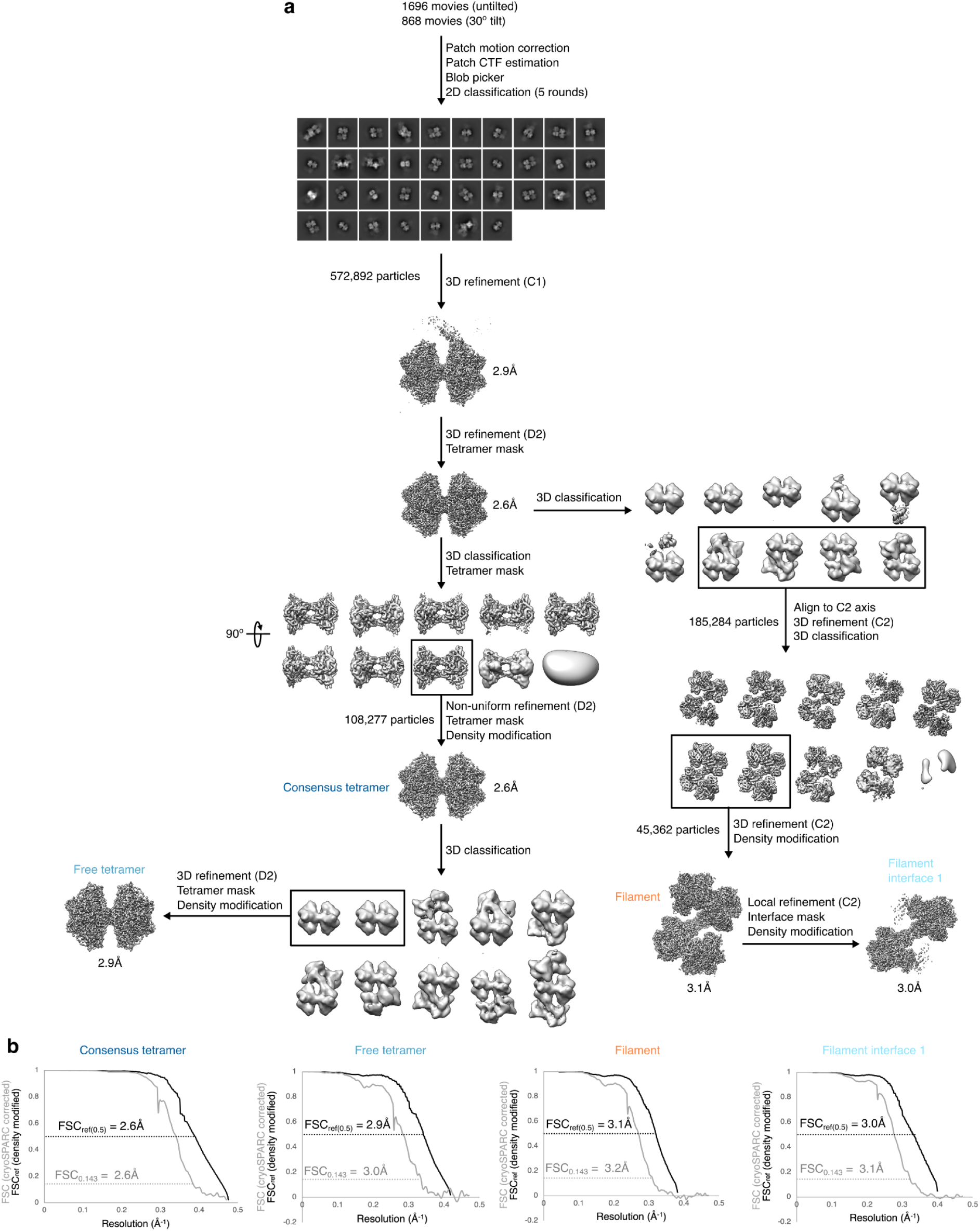
cryoEM data processing for T-state PFKL. **a,** Flowchart of cryoEM data processing. **b,** Noise-substituted corrected FSC curves (grey lines) and FSCref curves after density modification (black lines) and corresponding resolution estimates for the T-state consensus tetramer, free tetramer, filament, and filament interface 1 structures.

**Fig. S3:**
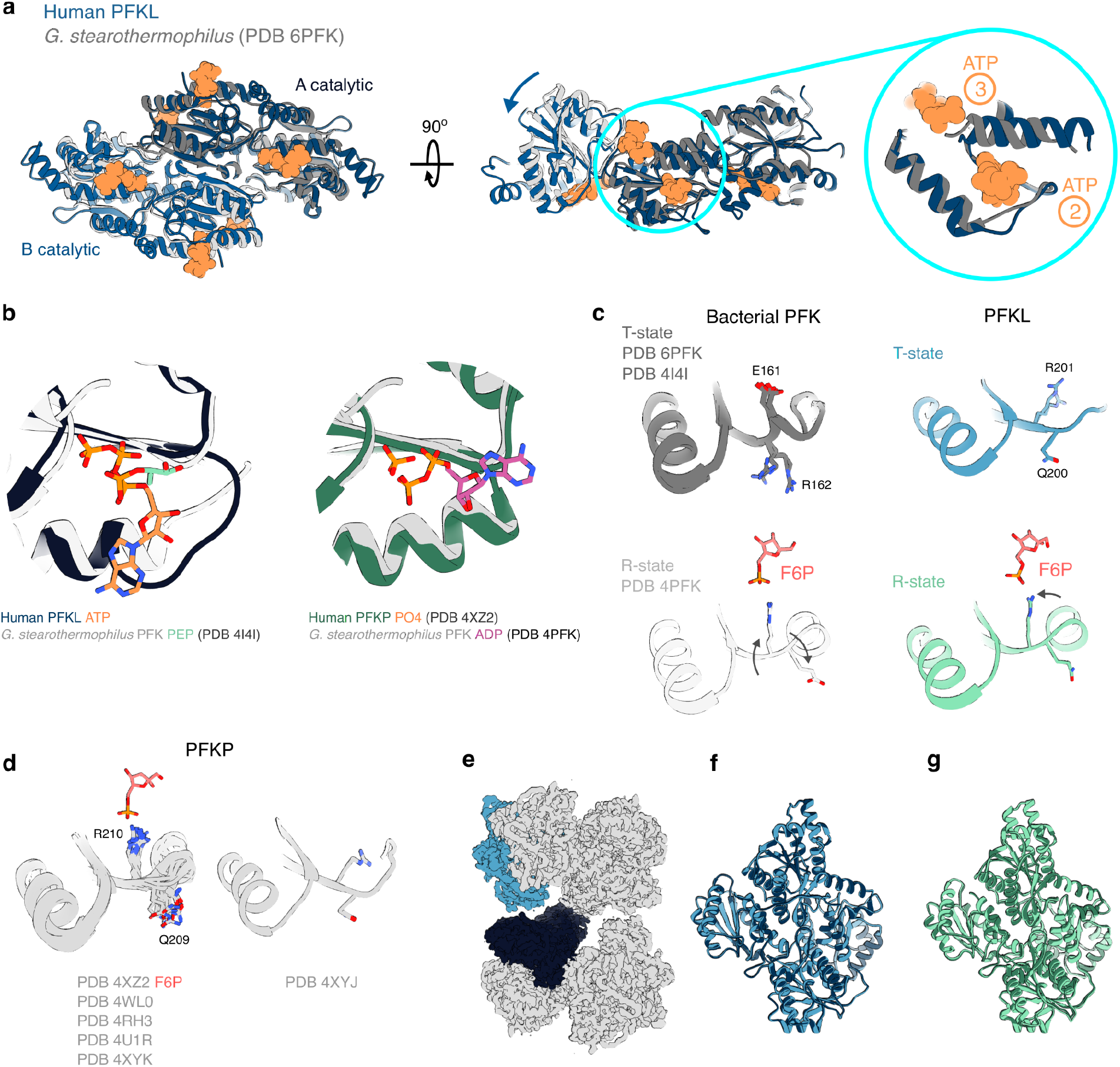
Conformational changes in PFKL. **a,** T-state Human PFKL and T-state bacterial PFK structures aligned on the catalytic domain of subunit A, with ATP in the human structure shown in orange. Human PFKL exhibits an additional rotation between adjoining catalytic domains (middle panel), as well as a shift in helix 376-389 adjacent to ATP sites 2 and 3 (expanded view in blue circle; right panel). **b,** ATP site 3 of PFKL is occupied by phosphoenolpyruvate (PEP) in T-state bacterial PFK (left) and by ADP in R-state bacterial PFK or phosphate in R-state PFKP (right). **c,** R162 and E161 in bacterial PFK swap orientations upon F6P binding and transition to the R-state, while the corresponding residues R201 and Q200 in PFKL show smaller rearrangements. R201 appears to occupy multiple positions in the cryoEM map. All chains from the indicated bacterial crystal structures are aligned and displayed. **d,** R210 is in the F6P-bound orientation in structures of PFKP (left), except for PDB 4XYJ (right). All chains from indicated crystal structures are aligned and displayed. F6P from PDB 4XZ2 is also shown. **e,** Cryo-EM structure of the T-state PFKL filament, with adjoining monomers at interface 2 colored in different shades of blue. **f,** Alignment of the monomers colored in **(e)**. **g,** Alignment of adjoining monomers at interface 2 from the R-state PFKL filament structure.

**Fig. S4:**
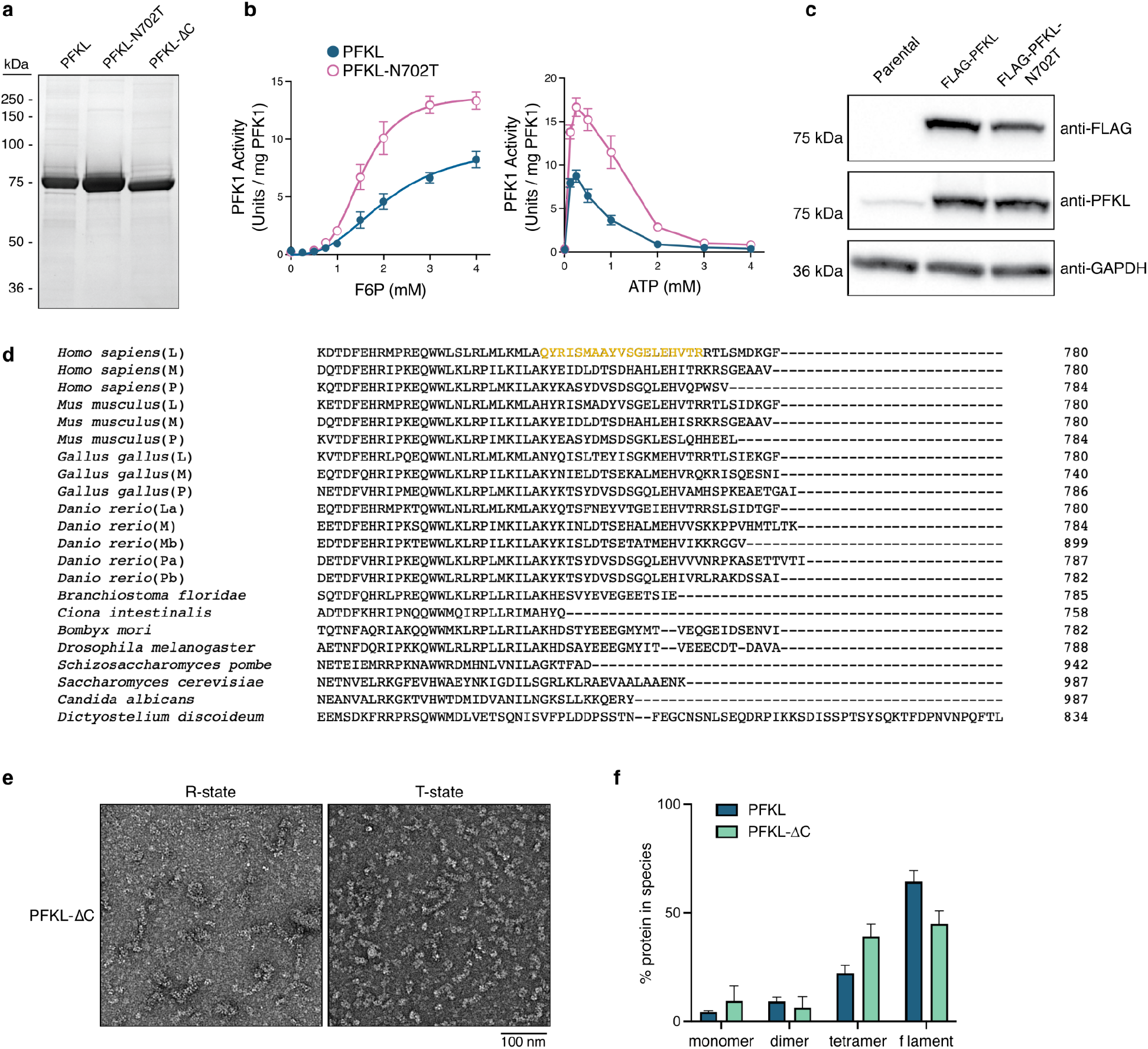
Characterization of wild type and mutant PFKL. **a,** Coomassie-stained SDS-PAGE gel with purified recombinant wild type and mutant PFKL. **b,** Enzyme activity assays to determine the affinity for F6P (left) and ATP (right) for wild type PFKL (blue) and PFKL-N702T (pink). Data for wild type PFKL are the same as shown in figure 4a and are reproduced here for comparison. Assay conditions are described in Table 1. Data are mean and SEM. **c,** Anti-FLAG and anti-PFKL western blots of parental, FLAG-PFKL, and FLAG-PFKL-N702T cells, with anti-GAPDH as a loading control. **d,** Sequence alignment of PFK1 C-termini from various species, with isoforms indicated as liver (L), muscle (M), or platelet (P) with subtypes as appropriate. The region of the PFKL C-terminus resolved in the T-state structure is indicated in yellow. **e,** Negative stain EM of PFKL-ΔC under R-state and T-state conditions. **f,** Quantification of assembly state of PFKL and PFKL-ΔC by mass photometry. Data for wild type PFKL are the same as shown in figure 3c and are reproduced here for comparison.

**Table S1.**
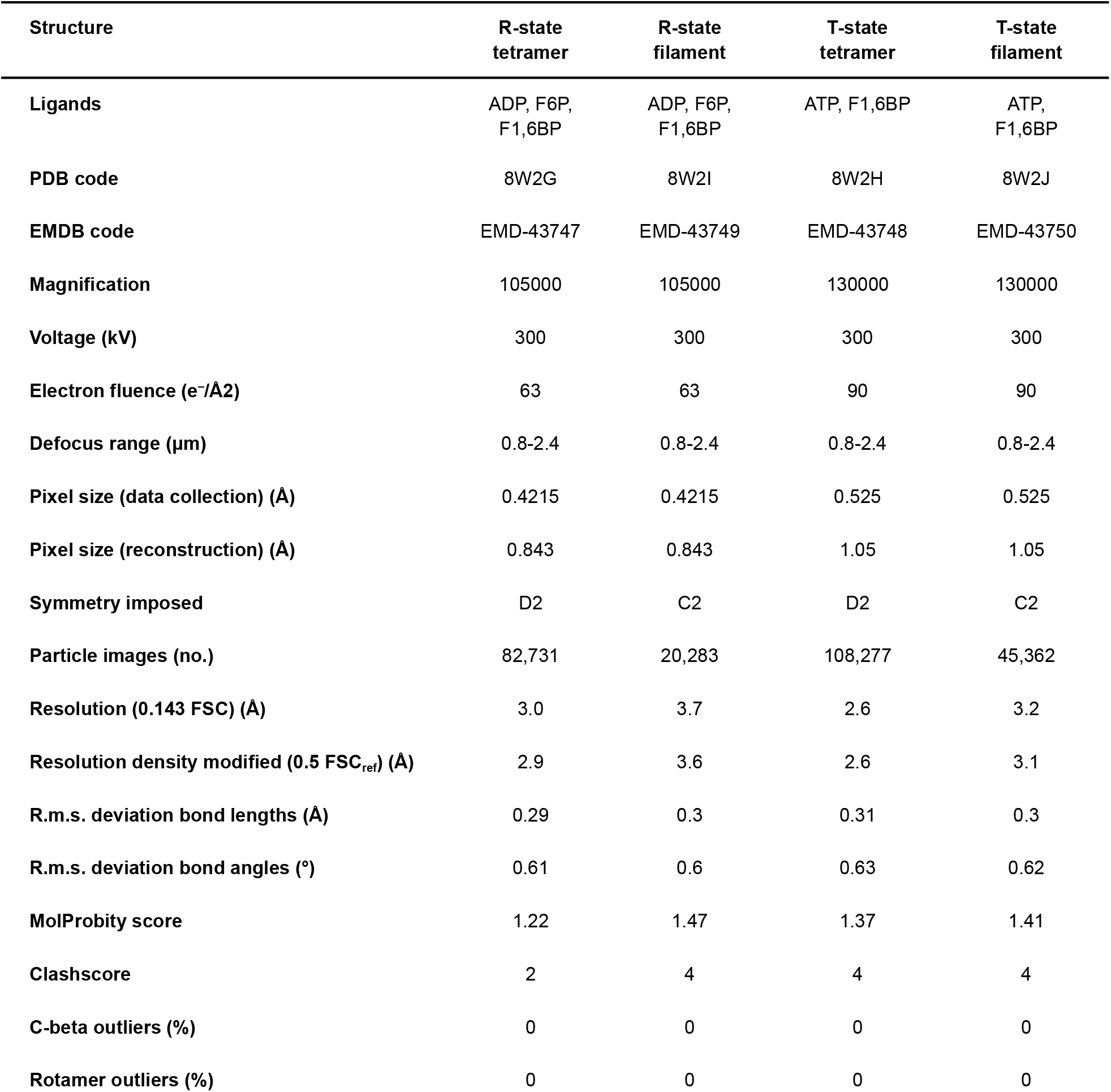
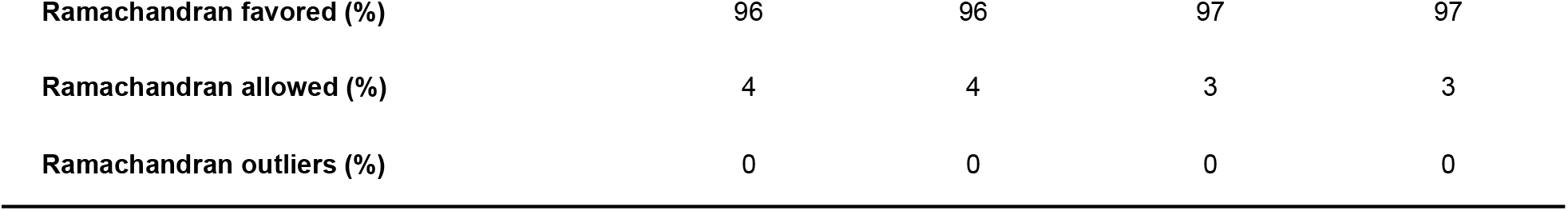
Cryo-EM data collection, refinement and validation statistics.

**Table S2:** Overview of the equilibration protocol for molecular dynamics simulations.

| Step | Length [ns] | $\Delta t$ [fs] | Restraints | $k_{BB}$ [kJ mol<sup>-1</sup> nm<sup>-2</sup>] | $k_{SC}$ [kJ mol<sup>-1</sup> nm<sup>-2</sup>] |
| --- | --- | --- | --- | --- | --- |
| 1 | 1.0 | 1 | heavy atoms | 1000 | 500 |
| 2 | 1.0 | 1 | heavy atoms | 500 | 250 |
| 3 | 1.0 | 1 | heavy atoms | 250 | 100 |
| 4 | 2.0 | 2 | heavy atoms | 100 | 50 |
| 5 | 2.0 | 2 | heavy atoms | 50 | 0 |
| 6 | 0.5 | 2 | none | 0 | 0 |

$k_{BB}$ - force constant of harmonic restraint potentials applied to the heavy atoms of the backbone and those of the ligands. $k_{SC}$ – force constant used to restrain side-chain heavy atoms. $\Delta t$ – time step.

## References

1. Evans, P. R., Farrants, G. W. & Lawrence, M. C. Crystallographic structure of allosterically inhibited phosphofructokinase at 7 A resolution. J. Mol. Biol. 191, 713–720 (1986).

2. Schirmer, T. & Evans, P. R. Structural basis of the allosteric behaviour of phosphofructokinase. Nature 343, 140–145 (1990).

3. Evans, P. R., Farrants, G. W. & Hudson, P. J. Phosphofructokinase: structure and control. Philos. Trans. R. Soc. Lond. B Biol. Sci. 293, 53–62 (1981).

4. Poorman, R. A., Randolph, A., Kemp, R. G. & Heinrikson, R. L. Evolution of phosphofructokinase—gene duplication and creation of new effector sites. Nature 309, 467–469 (1984).

5. Kemp, R. G. & Gunasekera, D. Evolution of the allosteric ligand sites of mammalian phosphofructo-1-kinase. Biochemistry 41, 9426–9430 (2002).

6. Banaszak, K. et al. The crystal structures of eukaryotic phosphofructokinases from baker’s yeast and rabbit skeletal muscle. J. Mol. Biol. 407, 284–297 (2011).

7. Webb, B. A. et al. Structures of human phosphofructokinase-1 and atomic basis of cancer-associated mutations. Nature 523, 111–114 (2015).

8. Kloos, M., Brüser, A., Kirchberger, J., Schöneberg, T. & Sträter, N. Crystal structure of human platelet phosphofructokinase-1 locked in an activated conformation. Biochem. J 469, 421–432 (2015).

9. Zancan, P., Marinho-Carvalho, M. M., Faber-Barata, J., Dellias, J. M. M. & Sola-Penna, M. ATP and fructose-2,6-bisphosphate regulate skeletal muscle 6-phosphofructo-1-kinase by altering its quaternary structure. IUBMB Life 60, 526–533 (2008).

10. Hesterberg, L. K. & Lee, J. C. Self-association of rabbit muscle phosphofructokinase: effects of ligands. Biochemistry 21, 216–222 (1982).

11. Costa Leite, T., Da Silva, D., Guimarães Coelho, R., Zancan, P. & Sola-Penna, M. Lactate favours the dissociation of skeletal muscle 6-phosphofructo-1-kinase tetramers down-regulating the enzyme and muscle glycolysis. Biochem. J 408, 123–130 (2007).

12. Hicks, K. G. et al. Protein-metabolite interactomics of carbohydrate metabolism reveal regulation of lactate dehydrogenase. Science 379, 996–1003 (2023).

13. Yi, W. et al. Phosphofructokinase 1 glycosylation regulates cell growth and metabolism. Science 337, 975–980 (2012).

14. Zhao, S. et al. Regulation of cellular metabolism by protein lysine acetylation. Science 327, 1000–1004 (2010).

15. Mahrenholz, A. M., Lan, L. & Mansour, T. E. Phosphorylation of heart phosphofructokinase by Ca2+ calmodulin protein kinase. Biochem. Biophys. Res. Commun. 174, 1255–1259 (1991).

16. Lee, J.-H. et al. Stabilization of phosphofructokinase 1 platelet isoform by AKT promotes tumorigenesis. Nat. Commun. 8, 949 (2017).

17. Fernandes, P. M., Kinkead, J., McNae, I., Michels, P. A. M. & Walkinshaw, M. D. Biochemical and transcript level differences between the three human phosphofructokinases show optimisation of each isoform for specific metabolic niches. Biochem. J 477, 4425–4441 (2020).

18. Webb, B. A., Dosey, A. M., Wittmann, T., Kollman, J. M. & Barber, D. L. The glycolytic enzyme phosphofructokinase-1 assembles into filaments. J. Cell Biol. 216, 2305–2313 (2017).

19. Amara, N. et al. Selective activation of PFKL suppresses the phagocytic oxidative burst. Cell 184, 4480–4494.e15 (2021).

20. Mosser, R., Reddy, M. C. M., Bruning, J. B., Sacchettini, J. C. & Reinhart, G. D. Redefining the role of the quaternary shift in Bacillus stearothermophilus phosphofructokinase. Biochemistry 52, 5421–5429 (2013).

21. Rizzo, S. C. & Eckel, R. E. Control of glycolysis in human erythrocytes by inorganic phosphate and sulfate. Am. J. Physiol. 211, 429–436 (1966).

22. Jin, M. et al. Glycolytic Enzymes Coalesce in G Bodies under Hypoxic Stress. Cell Rep. 20, 895–908 (2017).

23. Adams, A. G., Bulusu, R. K. M., Mukhitov, N., Mendoza-Cortes, J. L. & Roper, M. G. Online Measurement of Glucose Consumption from HepG2 Cells Using an Integrated Bioreactor and Enzymatic Assay. Anal. Chem. 91, 5184–5190 (2019).

24. Santamaria, B., Estevez, A. M., Martinez-Costa, O. H. & Aragon, J. J. Creation of an allosteric phosphofructokinase starting with a nonallosteric enzyme. The case of dictyostelium discoideum phosphofructokinase. J. Biol. Chem. 277, 1210–1216 (2002).

25. Huang, J. et al. CHARMM36m: an improved force field for folded and intrinsically disordered proteins. Nat. Methods 14, 71–73 (2017).

26. Yugi, K. et al. Reconstruction of insulin signal flow from phosphoproteome and metabolome data. Cell Rep. 8, 1171–1183 (2014).

27. Lynch, E. M., Kollman, J. M. & Webb, B. A. Filament formation by metabolic enzymes—A new twist on regulation. Curr. Opin. Cell Biol. 66, 28–33 (2020).

28. Simonet, J. C., Burrell, A. L., Kollman, J. M. & Peterson, J. R. Freedom of assembly: metabolic enzymes come together. Mol. Biol. Cell 31, 1201–1205 (2020).

29. Hvorecny, K. L. & Kollman, J. M. Greater than the sum of parts: Mechanisms of metabolic regulation by enzyme filaments. Curr. Opin. Struct. Biol. 79, 102530 (2023).

30. Garcia-Seisdedos, H., Empereur-Mot, C., Elad, N. & Levy, E. D. Proteins evolve on the edge of supramolecular self-assembly. Nature 548, 244–247 (2017).

31. Seisdedos, H. G., Levin, T., Shapira, G., Freud, S. & Levy, E. D. Mutant libraries reveal negative design shielding proteins from supramolecular self-assembly and relocalization in cells. Proceedings of the National Academy of Sciences vol. 119 Preprint at 10.1073/pnas.2101117119 (2022).

32. Hunkeler, M. et al. Structural basis for regulation of human acetyl-CoA carboxylase. Nature 558, 470–474 (2018).

33. Lynch, E. M. et al. Human CTP synthase filament structure reveals the active enzyme conformation. Nat. Struct. Mol. Biol. 24, 507–514 (2017).

34. Barry, R. M. et al. Large-scale filament formation inhibits the activity of CTP synthetase. Elife 3, e03638 (2014).

35. Stoddard, P. R. et al. Polymerization in the actin ATPase clan regulates hexokinase activity in yeast. Science 367, 1039–1042 (2020).

36. Lynch, E. M. & Kollman, J. M. Coupled structural transitions enable highly cooperative regulation of human CTPS2 filaments. Nat. Struct. Mol. Biol. 27, 42–48 (2020).

37. Pony, P., Rapisarda, C., Terradot, L., Marza, E. & Fronzes, R. Filamentation of the bacterial bi-functional alcohol/aldehyde dehydrogenase AdhE is essential for substrate channeling and enzymatic regulation. Nat. Commun. 11, 1426 (2020).

38. Kim, G. et al. Aldehyde-alcohol dehydrogenase undergoes structural transition to form extended spirosomes for substrate channeling. Commun Biol 3, 298 (2020).

39. Hu, H.-H. et al. Filamentation modulates allosteric regulation of PRPS. Elife 11, (2022).

40. Burrell, A. L. et al. IMPDH1 retinal variants control filament architecture to tune allosteric regulation. Nat. Struct. Mol. Biol. 29, 47–58 (2022).

41. Johnson, M. C. & Kollman, J. M. Cryo-EM structures demonstrate human IMPDH2 filament assembly tunes allosteric regulation. eLife vol. 9 Preprint at 10.7554/elife.53243 (2020).

42. Hvorecny, K. L., Hargett, K., Quispe, J. D. & Kollman, J. M. Human PRPS1 filaments stabilize allosteric sites to regulate activity. Nat. Struct. Mol. Biol. 30, 391–402 (2023).

43. Jang, S. et al. Glycolytic Enzymes Localize to Synapses under Energy Stress to Support Synaptic Function. Neuron 90, 278–291 (2016).

44. Kohnhorst, C. L. et al. Identification of a multienzyme complex for glucose metabolism in living cells. J. Biol. Chem. 292, 9191–9203 (2017).

45. Brüser, A., Kirchberger, J., Kloos, M., Sträter, N. & Schöneberg, T. Functional linkage of adenine nucleotide binding sites in mammalian muscle 6-phosphofructokinase. J. Biol. Chem. 287, 17546–17553 (2012).

46. Suloway, C. et al. Automated molecular microscopy: the new Leginon system. J. Struct. Biol. 151, 41–60 (2005).

47. Punjani, A., Rubinstein, J. L., Fleet, D. J. & Brubaker, M. A. cryoSPARC: algorithms for rapid unsupervised cryo-EM structure determination. Nat. Methods 14, 290–296 (2017).

48. Adams, P. D. et al. PHENIX: a comprehensive Python-based system for macromolecular structure solution. Acta Crystallogr. D Biol. Crystallogr. 66, 213–221 (2010).

49. Croll, T. I. ISOLDE: a physically realistic environment for model building into low-resolution electron-density maps. Acta Crystallogr D Struct Biol 74, 519–530 (2018).

50. Pettersen, E. F. et al. UCSF Chimera--a visualization system for exploratory research and analysis. J. Comput. Chem. 25, 1605–1612 (2004).

51. Young, G. et al. Quantitative mass imaging of single biological macromolecules. Science 360, 423–427 (2018).

52. Voronkova, M. A. et al. Cancer-associated somatic mutations in human phosphofructokinase-1 reveal a critical electrostatic interaction for allosteric regulation of enzyme activity. Biochem. J 480, 1411–1427 (2023).

53. The PyMOL Molecular Graphics System, Version 2.5.4 Schrödinger, LLC.

54. Abraham, M. J. et al. GROMACS: High performance molecular simulations through multi-level parallelism from laptops to supercomputers. SoftwareX 1-2, 19–25 (2015).

55. Jo, S., Kim, T., Iyer, V. G. & Im, W. CHARMM-GUI: a web-based graphical user interface for CHARMM. J. Comput. Chem. 29, 1859–1865 (2008).

56. Kim, S. et al. CHARMM-GUI ligand reader and modeler for CHARMM force field generation of small molecules. J. Comput. Chem. 38, 1879–1886 (2017).

57. Vanommeslaeghe, K. et al. CHARMM general force field: A force field for drug-like molecules compatible with the CHARMM all-atom additive biological force fields. J. Comput. Chem. 31, 671–690 (2010).

58. Jorgensen, W. L., Chandrasekhar, J. & Madura, J. D. Comparison of simple potential functions for simulating liquid water. The Journal of (1983).

59. Feenstra, K. A., Hess, B. & Berendsen, H. J. C. Improving efficiency of large time-scale molecular dynamics simulations of hydrogen-rich systems. J. Comput. Chem. 20, 786–798 (1999).

60. Darden, T., York, D. & Pedersen, L. Particle mesh Ewald: An N⋅log(N) method for Ewald sums in large systems. J. Chem. Phys. 98, 10089–10092 (1993).

61. Hockney, R. W., Goel, S. P. & Eastwood, J. W. Quiet high-resolution computer models of a plasma. J. Comput. Phys. 14, 148–158 (1974).

62. Hess, B., Bekker, H., Berendsen, H. J. C. & Fraaije, J. G. E. M. LINCS: A linear constraint solver for molecular simulations. J. Comput. Chem. 18, 1463–1472 (1997).

63. Miyamoto, S. & Kollman, P. A. Settle: An analytical version of the SHAKE and RATTLE algorithm for rigid water models. J. Comput. Chem. 13, 952–962 (1992).

64. Berendsen, H. J. C., van Postma, J., Van Gunsteren, W. F., DiNola, A. & Haak, J. R. Molecular dynamics with coupling to an external bath. J. Chem. Phys. 81, 3684–3690 (1984).

65. Bussi, G., Donadio, D. & Parrinello, M. Canonical sampling through velocity rescaling. J. Chem. Phys. 126, 014101 (2007).

66. Parrinello, M. & Rahman, A. Polymorphic transitions in single crystals: A new molecular dynamics method. J. Appl. Phys. 52, 7182–7190 (1981).

67. Vallat, R. Pingouin: statistics in Python. J. Open Source Softw. 3, 1026 (2018).

